# Chytrids-conveyed long-chain polyunsaturated fatty acids to Daphnia alleviate the detrimental effect of heat when combined with limiting dietary organic matter quantity and nutritional quality

**DOI:** 10.1101/2022.11.03.514985

**Authors:** András Abonyi, Matthias Pilecky, Serena Rasconi, Robert Ptacnik, Martin J. Kainz

## Abstract

Global warming enhances the dominance of poorly palatable PUFA-deprived bloom-forming cyanobacteria. Chytrid fungal parasites increase herbivory and dietary access to polyunsaturated fatty acids (PUFA) across the phytoplankton-zooplankton interface. Little is known however about the role chytrids may play in compensating for the decrease of algae-derived PUFA under global warming scenarios. We tested experimentally the combined effects of water temperature increase and the presence of chytrids with *Daphnia magna* as the consumer and the cyanobacterium *Planktothrix rubescens* as the main diet. We hypothesised that the diet including chytrids would enhance *Daphnia* fitness due to increased PUFA transfer irrespective of water temperature. Chytrid-infected diet significantly increased *Daphnia* survival, somatic growth, and reproduction, irrespective of water temperature. The PUFA content of *Daphnia* feeding on the chytrid-infected diet was unaffected by heat at the onset of the first successful reproduction. Carbon stable isotopes of fatty acids highlighted preferential n-3 PUFA upgrading by chytrids and an ~3x higher endogenous n-3 PUFA conversion compared with n-6 PUFA by *Daphnia*, irrespective of water temperature. Diet including chytrids enhanced the retention of eicosapentaenoic acid (EPA; 20:5n-3) and arachidonic acid (ARA; 20:4n-6) in *Daphnia*. The heat did not decrease EPA and even increased ARA retention by enhanced endogenous bioconversion in *Daphnia* when feeding on the chytrid-infected diet. We conclude that chytrids support *Daphnia* fitness at higher water temperatures via increased n-3 and n-6 PUFA retention and preferential n-3 PUFA bioconversion. Thus, they help function pelagic ecosystems with PUFA availability at the phytoplankton-zooplankton interface in a warmer climate.

## Introduction

Recent evidence suggests that chytrid fungal parasites perform key functions at the phytoplankton-zooplankton interface. Chytrids enhance herbivory via the fragmentation of poorly edible algal/cyanobacteria hosts (Frenken et al., 2020) and improve the dietary supply of polyunsaturated fatty acids (PUFA; Rasconi et al., 2020; Gerphagnon et al., 2019) and sterols (Kagami et al., 2007; Gerphagnon et al., 2019), as essential molecules, for zooplankton. Chytrids are ubiquitous invading various phytoplankton hosts (Gleason et al., 2015) and produce edible (~2-5 μm) free-swimming zoospores for subsequent consumers, such as zooplankton. Chytrids thus provide an alternative dietary energy pathway between phytoplankton and zooplankton, called the ‘mycoloop’ (Kagami et al., 2014). Chytrids upgrade dietary carbon from the algal/canobacteria hosts in terms of omega-3 PUFA and sterols (Gerphagnon et al., 2019; Rasconi et al., 2020), which positive dietary effect extends to *Daphnia* as a susequent concumer (Abonyi et al., 2022). Whether and how the mycoloop can support zooplankton with enhanced PUFA availability under global warming (IPCC, 2021), remains largely unexplored.

The efficient transfer of dietary energy and essential molecules across the phytoplankton-zooplankton trophic interface is crucial for consumers at higher trophic levels. The energy transfer can be constrained quantitatively and qualitatively. A quantitative limitation may occur in response to reduced primary production, e.g., during oligotrophication (Jeppesen et al., 2002; Jochimsen et al., 2013). Alternatively, increased primary production in response to eutrophication and global warming may result in the more frequent proliferation of deleterious phytoplankton (Paerl and Huisman, 2008; O’Neil et al., 2012). Bloom-forming cyanobacteria are typically filamentous and/or colonial, which hinders zooplankton grazing and creates an energetic bottleneck between pelagic primary producers and herbivorous consumers (Havens, 2008). Qualitative limitation of energy transfer may also be linked to phytoplankton blooms. Especially cyanobacteria lack essential compounds for zooplankton, such as PUFA and sterols, so their proliferation truncates trophic transfer of highly required dietary compounds.

Zooplankton cannot synthesize PUFA *de novo* (Gulati and Demott, 1997; Demott and Müller-Navarra, 1997). The omega-3 PUFA eicosapentaenoic acid (EPA, 20:5n-3) and the omega-6 PUFA arachidonic acid (ARA, 20:4n-6) are two conditionally indispensable molecules to support zooplankton growth and reproduction (see e.g. Sikora et al., 2016; Müller-Navarra et al., 2000; Ilić et al., 2021). They can be obtained directly from the diet or converted from their respective precursors linoleic acid (LIN, 18:2n-6) and alpha-linolenic acid (ALA, 18:3n-3), respectively. A key source of PUFA for herbivorous zooplankton in aquatic food webs is phytoplankton, which vary widely in PUFA composition (Taipale et al., 2013). Diatoms are rich in EPA, Cryptophytes contain EPA and docosahexaenoic acid (DHA, 22:6n-3), and thus both groups are considered high-quality diets for zooplankton. Green algae contain mostly LIN and ALA and are referred to as intermediate diet quality, while cyanobacteria generally lack long-chain PUFA and sterols (but see also the gamma-linoleic acid, GLA, 18:3n-6, content of *Microcystis* sp.; Strandberg et al., 2020), and are thus low-quality diets for zooplankton (Martin-Creuzburg et al., 2009).

Hardly palatable cyanobacteria of poor diet quality are deleterious for zooplankton (Bednarska et al., 2014), representing the worst-case scenario in trophic transfer of energy and essential dietary molecules. Recent evidence suggests that chytrid fungal parasites can support zooplankton fitness during cyanobacteria blooms (Agha et al., 2016; Abonyi et al., 2022). Diets including chytrids enhanced the somatic growth and reproduction of cladocerans (Kagami et al., 2007; Agha et al., 2016), copepods (Kagami et al., 2011), and rotifers (Frenken et al., 2018). Mechanisms potentially underlying the positive dietary effects of chytrids are 1) enhanced herbivory due to fragmentation of otherwise inedible cyanobacteria (Agha et al., 2016; Frenken et al., 2020), and; 2) increased nutritional quality due to PUFA conveyed or converted from the algal/cyanobacteria host (Gerphagnon et al., 2019; Taube et al., 2019; Rasconi et al., 2020). Chytrids can upgrade short-chain to long-chain PUFA from the host, i.e., convert the omega-3 PUFA ALA to stearidonic acid (SDA) and then further to long-chain PUFA, such as EPA (Rasconi et al., 2020). Recent evidence suggests that SDA synthesised by chytrids is selectively retained by *Daphnia* with a concomitant increase in the fitness of the consumer (Abonyi et al., 2022). How the mycoloop and the conveyed PUFA interact with heat reamains a key issue, especially for ectothermic pelagic consumers. Warming amplifies the deleterious effect of reduced diet intake on ectothermic organisms (Huey and Kingsolver, 2019). Accordingly, hardly edible PUFA-deprived cyanobacteria blooms may result in the collaps of herbivorous consumers, especially under water temperaute increase. In light of ongoing global warming (IPCC, 2021) and the more frequent occurrence of cyanobacteria blooms in aquatic ecosystems, it is crucial to test the strenght of the mycoloop to warmer conditions. If improved dietary PUFA provision via chytrids enhances *Daphnia* fitness on top of increased herbivory (Abonyi et al., 2022), it may also able to provide insurance against increasing water temperature.

Here we ask whether and how chytrids can support *Daphnia* fitness under water temperature increase and stress induced by the detrimental interaction of heat and hardly available dietary carbon and PUFA. Masclaux et al. (2009) showed a trade-off in *Daphnia* growth rate along with diet quality and water temperature. If high quality diet was available *ad libitum*, water temperature increase enhanced *Daphnia* growth, while reduced diet quality or quantity alleviated the positive temperature effect (Masclaux et al., 2009). On the other hand, multiple observations showed that *Daphnia* growth rate decreased at water temperatures >20°C (Giebelhausen and Lampert 2001; Masclaux et al., 2009) but it still remained a largely optimal condition for growth under non-limiting diet conditions (Müller et al., 2018). *Daphnia* adjust FA content to the water temperature, with decreasing PUFA towards higher temperatures (Zeis et al., 2019). Dietary EPA limitation was also shown to be more critical at lower than at higher water temperatures (Sperfeld and Wacker, 2012). The fitness of *Daphnia magna* is already constrained at 18°C when feeding on the poor diet quality and hardly edible *Planktothrix rubescens* (Abonyi et al., 2022). We thus performed a feeding experiment in which *Daphnia magna* was supplied with *Planktothrix rubescens* alone — a filamentous bloom-forming cyanobacterium — or with chytrid-infected *Planktothrix*. The two alternative diets were combined with two water temperature treatments, i.e. ambient (18°C) and heat (ambient +6°C). Applying a 24°C heat treatment enables us to study the PUFA transfer via the mycoloop with water temperature increase, without directly inducing *Daphnia* mortality by heat-mediated metabolic stress (Zeis et al., 2019; Im et al., 2020). We hypothesise that a +6°C water temperature increase in combination with a filamentous cyanobacterium diet would be detrimental to *Daphnia* fitness, also mediated by the PUFA-deprived diet conditions. We expect that diets including chytrids would support *Daphnia* fitness due to increased PUFA transfer even under increased water temperature. We examine survival, somatic growth, and reproduction of *Daphnia* as well as PUFA accrual of *Daphnia* among the treatments. In a ^13^C-labelling approach, we also aim at identifying the trophic carbon pathways via which physiologically functional PUFA are transferred to and bioconverted by *Daphnia*.

## Material and Methods

### Experimental cultures

The filamentous cyanobacterium *Planktothrix rubescens* (strain NIVA-CYA97/1) was cultured and maintained alone and together with its specific chytrid fungal parasite (strain Chy-Lys2009) on WC Medium (Guillard & Lorenzen, 1972). Cell culture bottles (VWR) were kept under non-axenic conditions at 21°C, applying 10.9 μmol m^2^ s^−1^ PAR and a 16:8 light:dark cycle in AquaLytic incubators (180 L, Liebherr, Germany). *Daphnia magna* was grown in pre-filtered (0.7 μm GF/F) lake water from Lake Lunz diluted with 10 V/V % of ADaM medium (Klüttgen et al., 1994). *Daphnia* were fed with 50% *Scenedesmus* sp. and 50% *Chlamydomonas* sp. >1 mg C L^−1^; both feeding cultures were also grown in WC medium.

### ^13^C-labelling of diet sources

*Planktothrix* and chytrid-infected *Planktothrix* diet cultures were labelled with ^13^C using NaH^13^CO_3_ (98 atom%, Sigma-Aldrich, USA) in the WC medium. Before each feeding occasion, diet cultures were diluted (1:1 v/v) with either non-labelled or labelled WC medium using SIMAX 2000 mL glass bottles filled up completely to avoid air space under the caps and kept under the culturing conditions for 24 hours. We did not manipulate water temperature for the diets, which would have modified their ^13^C values and made it thus impossible to compare *Daphnia* response among the treatments to diet *versus* heat. We expressed the ^13^C uptake efficiency based on all the pairwise differences in δ^13^C_FA_ between the labelled and the non-labelled replicates for each FA (i.e. Δ^13^C_FA_ = δ^13^C_FA-labelled_-δ^13^C_non-labelled_).

### Experimental design

The two diet treatments were: 1) *Planktothrix* (cyanobacterium—’CY’), and; 2) chytrid-infected *Planktothrix* (cyanobacterium-parasite—’CYP’), containing also free-swimming chytrid zoospores (zoospores—’Z’). Two water temperature treatments were applied: 1) keeping the experimental bottles at 18°C (temperature set in the experimental climate room), or; 2) keeping the experimental bottles in a 24±1°C water bath using a JBL ProTemp S 25 (Germany) heater. The water temperature in the experimental bottles (1 L Corning® Square Storage) varied by ±0.5°C during the entire experiment. The *Daphnia* were either fed with non-labelled or with ^13^C-labelled cultures. All treatments – applying a full factorial design – were performed in triplicates, resulting in 24 experimental units.

At the start of the experiment, bottles contained 30 *Daphnia* neonates that were <48 hrs old. The volume was constantly kept at 1 L, and the bottles were covered with a non-transparent plastic plate in a non-hermetic way. A 16:8 light:dark cycle with 1.4 μmol m^2^ s^−1^ PAR during the day-light phase was applied. We replaced the medium and provided new diets to *Daphnia* every other day corresponding to 1±0.01 μg C L^−1^ in all treatments. To do so, first the carbon content of the cultures were analysed prior to the experiment (Abonyi et al., 2022), and then, we approximated the carbon based on dry-weight per volume for each feeding occasion (i.e., adding different diet volumes to meet the 1μg C L^−1^). *Daphnia* were observed daily for survival and egg production. Individuals lying at the bottom of the bottles with no movement for 5 minutes were considered dead and removed. To reliably compare the PUFA content of *Daphnia* across treatments, ambient and heat treatments were stopped separately at the onset of the first successful hatching.

### *Life history of* Daphnia

The somatic growth rate (GR) of *Daphnia* was assessed as GR = (ln(DW_end_) - ln(DW_start_)) / d, where *DW*_*end*_ was the average individual dry weight in each experimental bottle at the end of the experiment, *DW*_*start*_ was the average individual dry weight of *Daphnia* neonates at the start of the experiment (average of 3 × 20 individuals), and *d* was the duration of the experiment in days. *Daphnia* survival was expressed as the % of surviving individuals. Egg production was quantified based on the number of eggs produced by *Daphnia* individuals and therefore standardised by the number of survivals (ind^−1^).

### *Elemental C, N, P compositions of diet and* Daphnia

The elemental C, N and P contents of diets and chytrid zoospores were analysed in triplicates at the beginning of the experiment (day 1), every second time of feeding (day 4, day 8, day 12) and right after the experiment (n=15). *Daphnia* were analysed for C, N, and P in triplicates at the start (T_0_) and at the end of the experiment (T_E_). We separated chytrid zoospores from *Planktothrix* filaments by filtering ~100 mL culture five times through a 20 μm mesh followed by a single filtration through a 5 μm mesh. The absence of *Planktothrix* in chytrid zoospore samples was evaluated using a Nikon Eclipse TS100 inverted microscope at 200 x magnification. Diets were collected on muffled and pre-weighed GF/F Whatman™ filters, dried for 48 hours, weighed and folded in tin capsules. *Daphnia* were first frozen (−80 °C), then freeze-dried for 48 hours (VirTis benchtopK, VWR), weighed, and put in tin capsules. C and N were measured using a Thermo Fisher Scientific Flash 2000 Elemental Analyser, linked to a Delta V Advantage Mass Spectrometer (for details, *see* Abonyi et al., 2022). The δ^13^C values were referenced to Vienna PeeDee Belemite (^13^C:^12^C = 0.01118) using USGS standards. P was measured as particulate organic phosphorus based on the ascorbic acid colourimetric method following persulfate digestion (APHA, 1998). Elemental C:N, C:P and N:P ratios were expressed as molar ratios.

### Fatty acids and compound-specific carbon isotope analyses

The fatty acids (FA) and their compound-specific carbon isotopes (i.e. CSI data on *δ*^13^C_FA_) were analysed in the diets (i.e., *Planktothrix*, chytrid-infected *Planktothrix* excluding chytrid zoospores, chytrid zoospores) and *Daphnia*. Individual diet sources were collected on muffled and pre-weighed GF/F Whatman™ filters (47 mm, 0.7 μm pore size), stored at −80°C then freeze-dried (VirTis benchtopK, VWR). *Daphnia* were also freeze-dried and weighted ~1 mg dry-weight into tin-caps. Lipids were extracted from the freeze-dried samples using a mix of chloroform-methanol (2:1), following the protocol described in Heissenberger et al. (2010).

Fatty acids were derivatised to methyl esters by incubation with 1% H_2_SO_4_ in methanol at 50 °C for 16 hours. Fatty acid methyl esters (FAME) were dried under N_2_ and dissolved in hexane. For quantification via flame ionisation, FAME were separated using a GC (Trace™ 1310 Thermo Specific, Italy) equipped with a Supelco™ SP-2560 column (100 m x 0.25 mm x 0.2 μm). ^13^C CSI analysis was performed using a Thermo Trace 1310 GC (ThermoFisher Scientific), connected via a ConFlo IV (Thermo Co.) to an Isotope Ratio Mass Spectrometer (IRMS, DELTA V Advantage, Thermo Co.). Samples were run against certified Me-C20:0 standards (USGS70: δ^13^C□=□−30.53‰, USGS71: δ^13^C□=□−10.5‰, and USGS72: δ^13^C□=□−1.54‰), which were used for drift and linear correction (for details, see Abonyi et al., 2022).

### *Quantitative PUFA retention by* Daphnia

We compared the retention of LIN, GLA, ARA, ALA, SDA, and EPA by *Daphnia* among treatments in two complementary ways: first, by comparing FA mass fractions per unit biomass (i.e. μg FA mg^−1^), and, second, based on the pairwise difference in δ^13^C_FA_ values between the labelled and the non-labelled replicates for each FA (i.e. Δ^13^C_FA_ = δ^13^C_FA-labelled_-δ^13^C_non-labelled_). The higher the value, the more efficient the PUFA retention, including the sources of dietary uptake and endogenous bioconversion. Furthermore, for comparing the efficiency of the endogenous conversion of short-chain (LIN, ALA) to long-chain PUFA (ARA, EPA) by *Daphnia*, we expressed the ratios of FA retentions between the pairs, i.e. Δ^13^C_ARA_ / Δ^13^C_LIN_ and Δ^13^C_EPA_ / Δ^13^C_ALA_. The higher the value, the more efficient the endogenous FA conversation.

### Statistical analyses

We tested for significant differences in fitness parameters (i.e. survival, somatic growth rates, eggs.ind^−1^,egg production rate), and PUFA retention efficiencies among treatments using Kruskal-Wallis rank sum tests (Hollander & Wolfe, 1973). In case of significant differences among the treatments, we ran a pairwise comparison of the treatment groups using the Wilcoxon rank sum test (Bauer, 1972). We controlled p values for multiple testing by applying the false discovery rate approach (Benjamini and Hochberg, 1995). We tested for significant differences in FA content between two groups (ambient and heat) using the Wilcoxon rank sum test. The FA data were ln(x+1) transformed for data visualisation and statistical analyses. The level of statistical significance was set at p<0.05 in all cases.

## Results

### Daphnia *fitness*

*Daphnia* feeding on the chytrid-infected diet under heat reproduced successfully on day 11, i.e. the end of the heat treatment. *Daphnia* on the ambient water temperature reproduced on day 15, i.e. the end of the ambient water temperature treatment. *Daphnia* feeding on the sole *Planktothrix* diet did not develop eggs during the experiment and started to die off after 3 days (Fig 1 AB). Their survival declined linearly under ambient water temperature but it did exponentially under heat.

**Fig 1.**
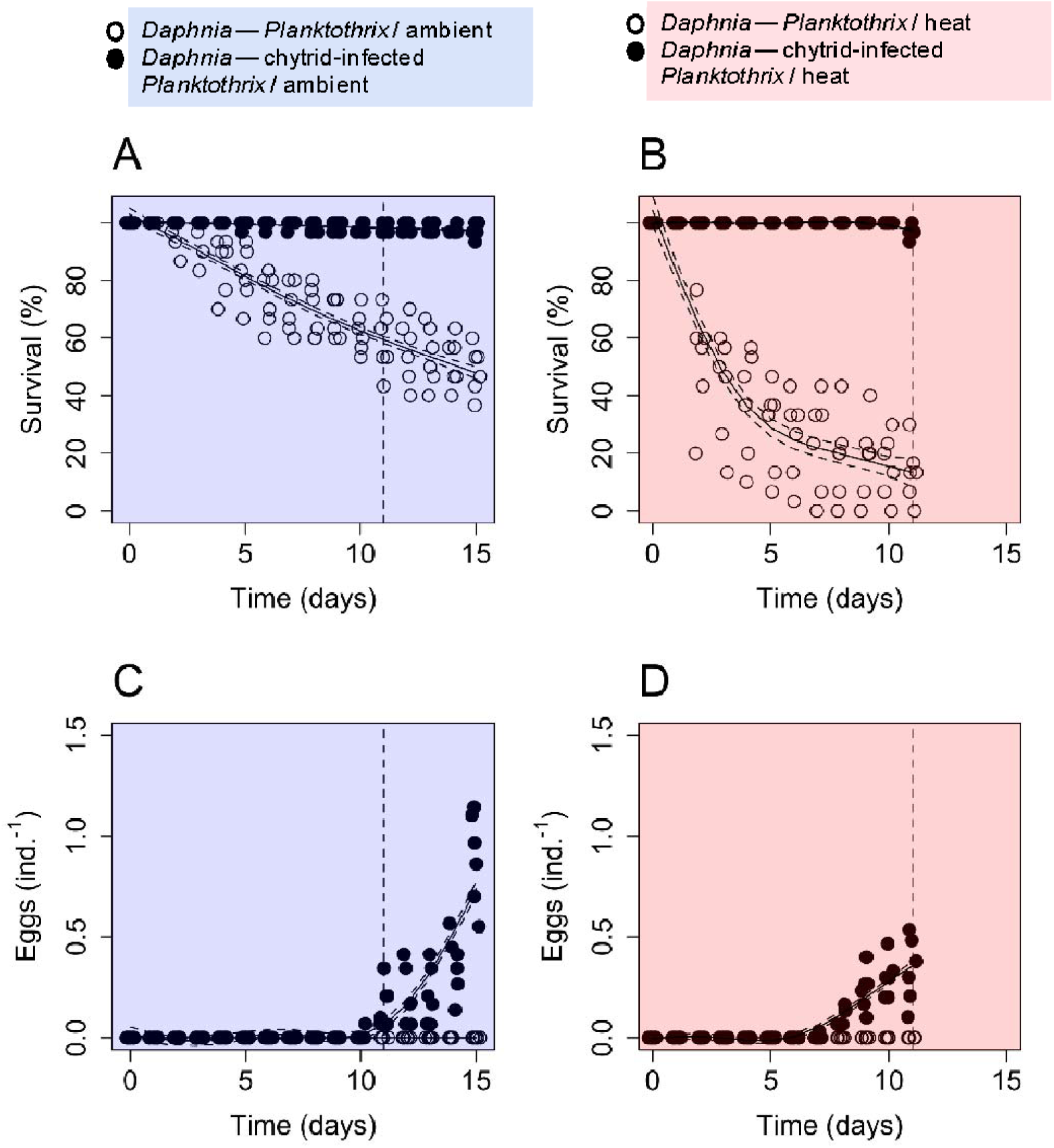
**(A)** *Daphnia* survival (% from 30 individuals per bottle) at ambient water temperature (18°C); and **(B)** *Daphnia* survival in the heat water temperature treatment (24°C); **(C)** the number of eggs produced per *Daphnia* at 18°C; and **(D)** the number of eggs produced per *Daphnia* at 24°C. Empty circles are *Daphnia* feeding on *Planktothrix*, while full circles represent *Daphnia* feeding on chytrid-infected *Planktothrix*. The vertical dashed lines indicate day 11, the onset of the first successful reproduction of *Daphnia* feeding on the chytrid-infected diet under heat (i.e. when the heat treatment was stopped).

Survival differed significantly among the treatment at day 11 (S1A, Kruskal-Wallis, chi-squared=19.751, df = 3, p<0.001). Heat significantly reduced survival only in Daphnia feeding on the sole Planktothrix diet (ambient: 58.9±10.9% (mean±SD), heat: 13.3±10.1%; pairwise Wilcoxon, p<0.01), while survival of Daphnia feeding on chytrid-infected diet remained unaffected (ambient: 97.8±1.7%, heat: 96.7±2.1%; pairwise Wilcoxon, p>0.05).

Somatic growth rates differed significantly among the treatments (S1B, Kruskal-Wallis, chi-squared = 17.561, df = 3, p<0.001). Somatic growth remained unaffected by heat (pairwise Wilcoxon, p>0.05) but was significantly enhanced by diet including chytrids (p<0.01).

*Daphnia* feeding on chytrid-infected diet started to produce eggs on day 7 under heat, while at the ambient water temperature onday 10 (FIG 1C,D). Heat reduced the number of eggs produced per *Daphnia* when compared at the day of the first successful reproduction (day 11 under heat: 0.33±0.16 eggs.ind-1 (mean±SD); day 15 under ambient condition: 0.89±0.23 eggs.ind^−1^ (Wilcoxon, W = 36, p<0.01).

Egg production rate differed significantly among the treatments (S1C, Kruskal-Wallis, chi-squared = 20.341, df = 3, p<0.001). It significantly increased in *Daphnia* feeding on chytrid-infected *Planktothrix* (pairwise Wilcoxon, p<0.01), which positive effect was significantly reduced by heat (pairwise Wilcoxon, p<0.01).

### *Individual PUFA transfer to* Daphnia

Due to the high mortality of *Daphnia* feeding on the sole *Planktothrix* diet, it was only possible to analyse the PUFA content in *Daphnia* feeding on the chytrid-infected diet. The individual PUFA contents in *Daphnia* feeding on chytrid-infected *Planktothrix* did not differ significantly between the ambient and heat treatments (Wilcoxon rank sum tests, p>0.05 in all cases, Fig 2). ALA had the highest mass fractions of the identified short-chain PUFA in *Daphnia* (ambient: 11.2±8.7 μg mg^−1^(mean±SD), heat: 6.0±0.6 μg mg^−1^) followed by LIN (ambient: 2.0±1.5 μg mg^−1^, heat: 1.5±0.1 μg mg^−1^) and SDA (ambient: 1.2±0.9 μg mg^−1^, heat: 0.7±0.1 μg mg^−1^) and GLA (ambient: 0.1± 0.08 μg mg^−1^, heat: 0.1±0.02 μg mg^−1^). EPA had the highest mass fractions of the identified long-chain PUFA in *Daphnia* (ambient: 0.9±0.5 μg mg^−1^, heat: 1.3 ±0.2 μg mg^−1^), followed by ARA (ambient: 0.5±0.3 μg mg^−1^, heat: 0.8± 0.1 μg mg^−1^) and DHA (ambient: 0.1±0.07 μg mg^−1^, heat: 0.05± 0.01 μg mg^−1^).

**Fig 2.**
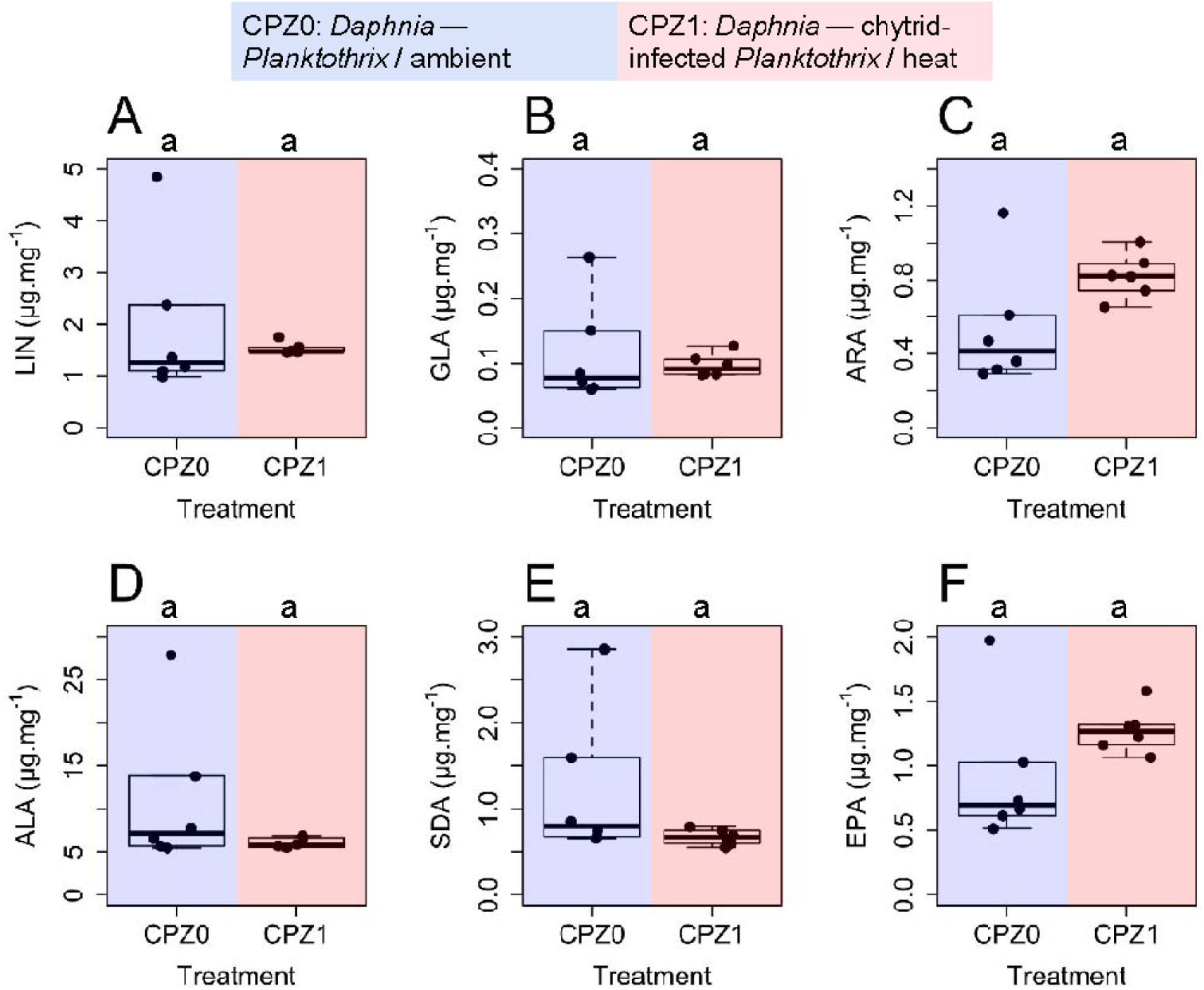
Polyunsaturated fatty acid (PUFA) content of *Daphnia* feeding on the chytrid-infected cyanobacterium *Planktothrix* and chytrid zoospores (CPZ) at the time of the first successful reproduction. CPZ0 stands for ambient water temperature (blue), while APZ1 stands for heat (red). **(A)** linoleic acid (18:2n-6, LIN); **(B)** gamma-linoleic acid (18:3n-6, GLA); **(C)** arachidonic acid (20:4n-6, ARA); **(D)** alpha-linolenic acid (18:3n-3, ALA); **(E)** stearidonic acid (18:4n-3, SDA); **(F)** eicosapentaenoic acid (20:5n-3, EPA). Letters denote the significance level for comparing values between water temperature treatments based on Wilcoxon rank sum tests (n=6 in all cases).

### ^*13*^*C-labelling of dietary PUFA and their retention by* Daphnia *based on CSI analysis*

The ^13^C uptake via LIN, ALA and SDA differed significantly among diet sources (Kruskal-Wallis rank sum tests, chi-squaredLIN = 415.27 (df=2), chi-squaredALA = 277.47 (df=2), chi-squaredSDA =295.48 (df=1), p<0.001 in all cases). ^13^C uptake via LIN was the highest in *Planktothrix* (438±229 ‰), significantly lower in chytrid-infected *Planktothrix* (284±104 ‰), and again, significantly lower in chytrid zoospores (34±26 ‰). ^13^C labelling of ALA was the highest in chytrid-infected *Planktothrix* (139±57 ‰), significantly lower in the cyanobacterium diet (121±60 ‰), and chytrid zoospores (43±33 ‰). SDA was absent in *Planktothrix* but present in the chytrid-infected cyanobacterium and the chytrid zoospores. ^13^C uptake via SDA was significantly higher in chytrid-infected *Planktothrix* (56±39 ‰) than in the chytrid zoospores (6±5 ‰). Diet sources did not contain ARA or APA (see also S3A).

Due to the high mortality of *Daphnia* feeding only on *Planktothrix* under heat, *Daphnia* from this treatment could not be analysed for FA. Among the other treatments, *Daphnia* significantly differed in LIN, ALA, SDA, ARA and EPA (Kruskal-Wallis rank sum tests, p_LIN_<0.01, p_ALA_<0.001, p_SDA_<0.001, p_ARA_<0.001, p_EPA_<0.01). At ambient water temperature, ^13^C-labelling indicated significantly enhanced LIN, ALA, SDA, ARA, and EPA retention in *Daphnia* feeding on the chytrid-infected *Planktothrix* diet (Pairwise Wilcoxon rank sum tests, p_LIN_<0.001, p_ALA_<0.01, p_SDA_<0.001, p_ARA_<0.001, p_EPA_<0.05; Fig 3). The heat treatment did not affect PUFA retention significantly in *Daphnia* feeding on the chytrid-infected diet (Pairwise Wilcoxon rank sum tests, p>0.05, in all cases), except ARA, where heat significantly enhanced its retention (Fig 3D, Pairwise Wilcoxon rank sum tests, p<0.01).

**Fig 3.**
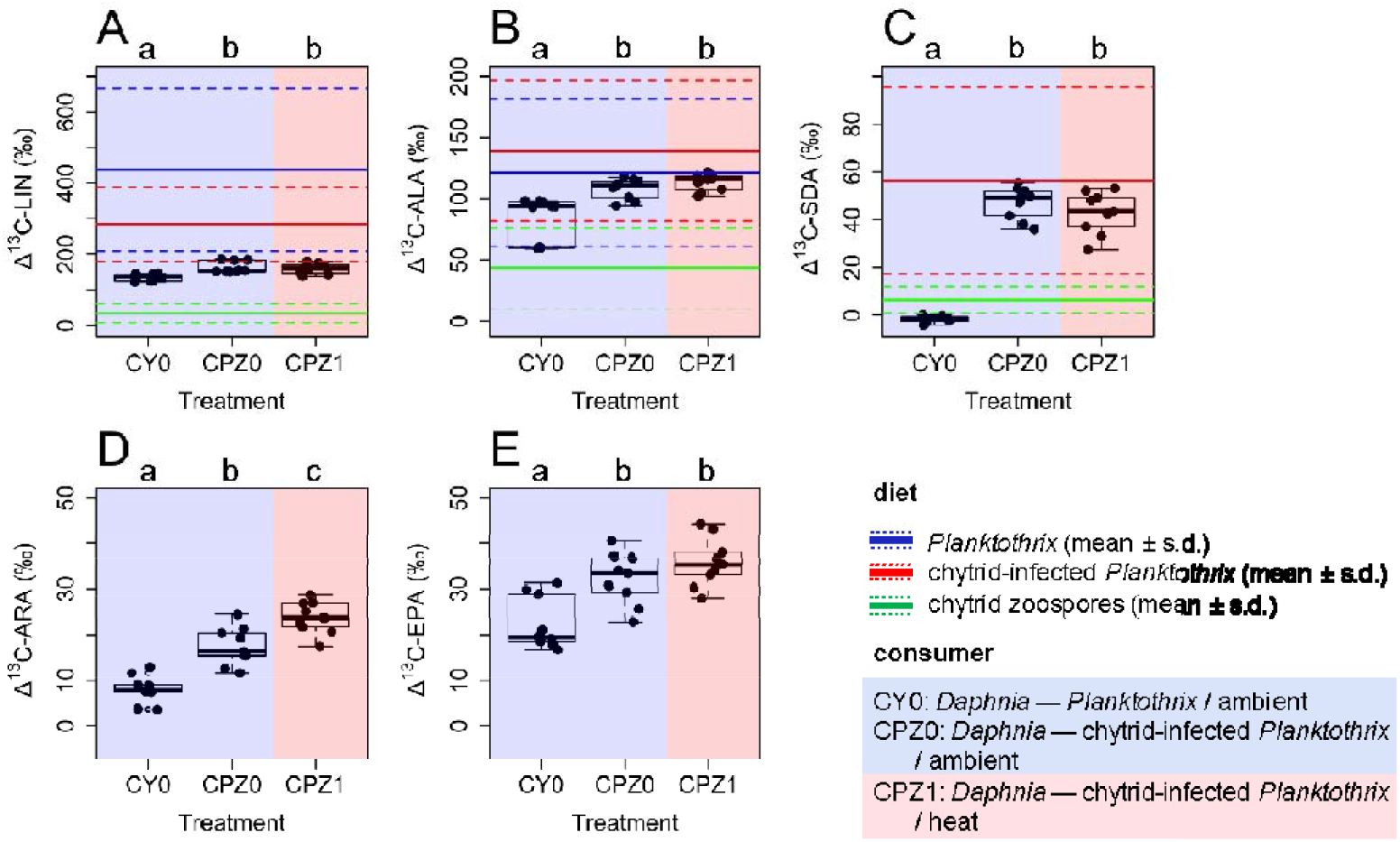
^13^C uptake via key polyunsaturated fatty acids (PUFA) in 1) the diet sources (lines): *Planktothrix* (blue), chytrid-infected *Planktothrix* (red) and chytrid zoospores (green) (all pairwise differences between labelled and non-labelled FA, n=225 for each pair) and 2) their retention by *Daphnia* (boxplots and raw data) at the onset of the first successful reproduction among treatments (all pairwise differences between labelled and non-labelled FA, n=9 for each pair) **(A)** linoleic acid (18:2n-6, LIN); **(B)** alpha-linolenic acid (18:3n-3, ALA); **(C)** stearidonic acid (18:4n-3, SDA); **(D)** arachidonic acid (20:4n-6, ARA); and **(E)** eicosapentaenoic acid (20:5n-3, EPA). Letters above the figures denote significant differences in FA retention of *Daphnia* among treatments (Pairwise Wilcoxon rank sum tests)

Endogenous conversion of short-chain to long-chain PUFA along with the n-3 chain (i.e. ALA to EPA) occurred with a ~3x higher conversion efficiency compared with the n-6 chain (i.e. LIN to ARA, Fig 4). The chytrid-infected diet alone and also in combination with heat significantly increased the LIN to ARA conversion (Fig 4A, Pairwise Wilcoxon rank sum tests, p<0.001, in all cases). Along with the ALA to EPA chain, the chytrid diet alone slightly but significantly (p<0.05), and the chytrid diet together with the heat highly significantly (p<0.001) enhanced the endogenous conversion (Fig 4B, Pairwise Wilcoxon rank sum tests).

**Fig 4.**
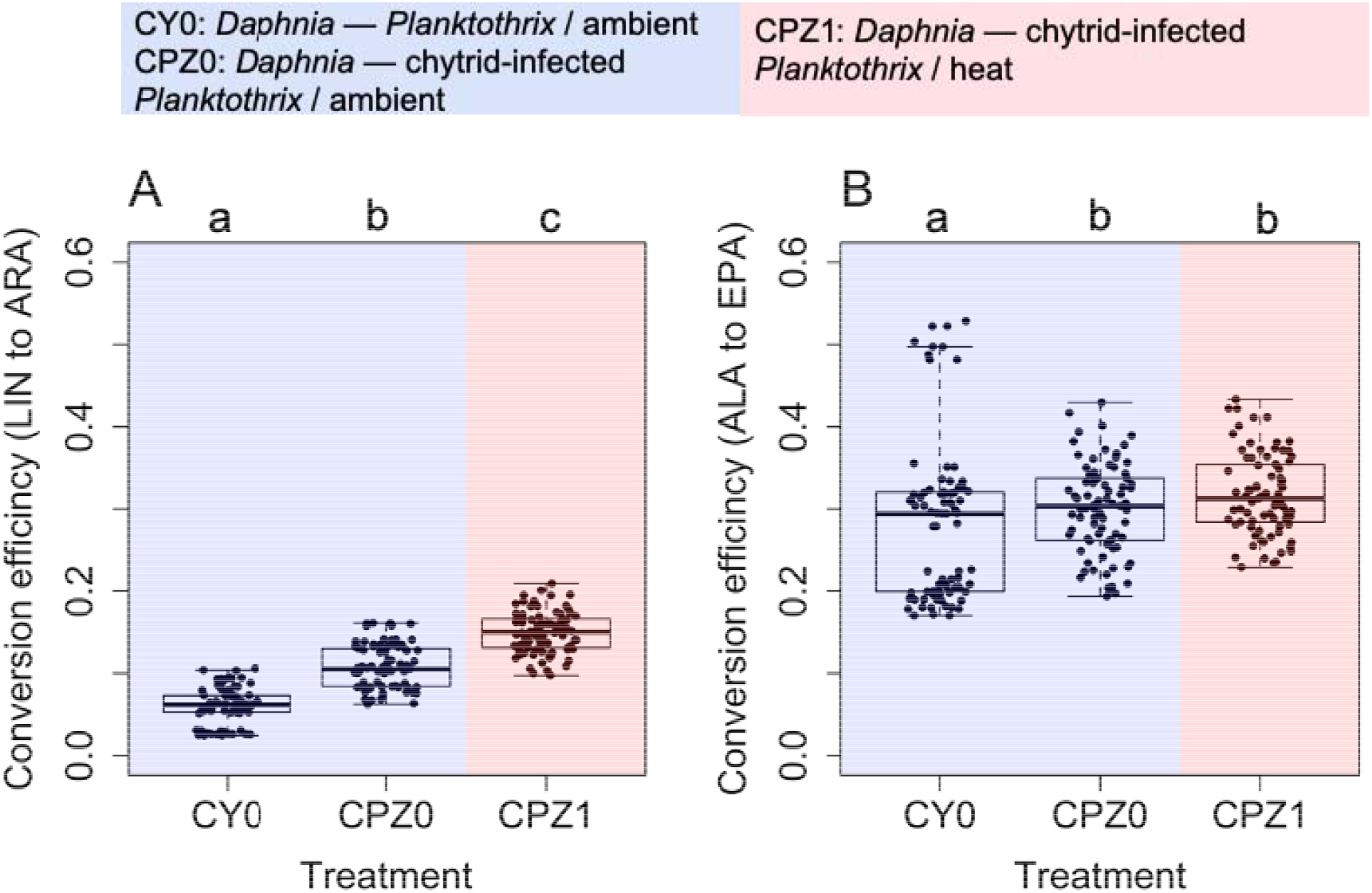
Efficiency in the endogenous bioconversion of **(A)** LIN to ARA (n-6 chain); and **(B)** ALA to EPA (n-3 chain) in *Daphnia* at the onset of the first successful reproduction among treatments (boxplots show all pairwise ratios between Δ^13^C_ARA_/Δ^13^C_LIN_ and Δ^13^C_EPA_/Δ^13^C_ALA_; n=81, for each treatment). Letters above the figures denote the level of significance in the difference in FA conversion of *Daphnia* among treatments (Pairwise Wilcoxon rank sum tests).

## Discussion

The high mortality of *Daphnia* neonates exposed to heat and the sole *Planktothrix* diet after after three days confirmed the hypothesis that a +6°C water temperature increase would be detrimental to the zooplankton consumer in case of diet limitation. Our data also confirmed that chytrid-infected diet could support *Daphnia* fitness under stress conditions induced by the combined impact of heat and limited diet availability. Chytrids could enhance the dietary supply and subsequent retention of key PUFA by *Daphnia* at both water temperature treatments. Our study thus provides a first experimental evidence for the positive diet quality effect of chytrids in terms of PUFA, as a possible key mechanism that may provide insurance against the detrimental effect of heat under cyanobacteria blooms in nature.

### PUFA transfer in the mycoloop under heat

Chytrids can perform trophic upgrading along with both the n-3 and n-6 PUFA chain (Rasconi et al., 2020), resulting in enhanced PUFA retention in Daphnia (Abonyi et al., 2022). Here we show that enhanced PUFA retention by *Daphnia* with chytrid-infected diet remains largely unaffected by a +6°C water temperature increase, if the diet conditions are limiting in terms of both organic matter quantity and nutritional quality. Due to the lack of ARA and EPA in the diet sources and the quantitative limitation in diet, LIN and ALA are converted into ARA and EPA by the *Daphnia* itself, respectively (Bec et al., 2003; Abonyi et al. 2022). The compound-specific fatty acid data clearly show that the endogenous bioconversion of ALA to EPA is more efficient than the conversion of LIN to ARA in *Daphnia*. While the conversion rate is assumed to be very low and the EPA content produced may still limit somatic growth (Ilić et al., 2021 and herein references), here we see that it is largely enhanced by chytrids, providing a plausible mechanistic link between chytrid-mediated enhanced n-3 PUFA retention and *Daphnia* fitness. The n-3 conversion pathway producing EPA, a key functional LC-FA for growth and reproduction (Becker & Boersma, 2003), remains unaffected by heat if chytrids are present in the diet. The sustained retention under heat also holds true for the n-3 PUFA SDA, as yet, however, with unknown energetic benefits (Abonyi et al., 2022). Surprisingly, while the chytrid diet enhances n-3 over n-6 PUFA retention (i.e. in response to enhanced endogenous conversion), the heat seems to significantly enhance the n-6 PUFA ARA retention via a more efficient LIN to ARA conversion. While this may suggest a crucial physiological role for ARA, its benefits under heat conditions remain to be further elucidated. The more efficient endogenous n-3 PUFA conversion and more stable EPA content in *Daphnia* may show that EPA is the limiting factor over ARA. Accordingly, as soon as a certain amount of EPA is acquired, *Daphnia* may reduce its conversion rate and start reproducing (Pilecky et al., 2021). This might be supported by the fact that EPA is preferentially stored by *Daphnia magna* and it appears in higher concentrations than ARA in the eggs (Becker and Boersma, 2005). If this is the case, high conversion rates from LIN to ARA might actually be an unnecessary - by-product - process for the animal, since, in case of its high need, ARA levels would also remain stable (Pilecky et al., 2021). On dietary PUFA, however, a recent study suggested that *Daphnia* could be even more susceptible to the lack of ARA compared with EPA (Ilić et al., 2021). As both ARA and EPA were lacking in diets, our results highlighting a preferential n-3 PUFA over n-6 PUFA conversion may not support the view of Ilic et al. (2021). Moreover, the potential effect of the chytrid-synthesised SDA on the endogenous conversion pathway of ALA to SDA still remains unknown. It is possible that the preferential n-3 PUFA conversion by the *Daphnia* might potentially have been enhanced by the favoured n-3 PUFA upgrading of chytrids (Rasconi et al., 2020; Abonyi et al., 2022), which would provide a mechanistic link between increased dietary quality for *Daphnia* and the enhanced performance of the animals.

### The chytrid-mediated PUFA insurance against the heat

Ectothermic organisms can genetically adapt to higher temperatures (Geerts et al., 2015). Also, the physiological plasticity of animals allows changes in biochemical characteristics to counter the negative impact of heat, for example, *Daphnia* adjust its FA content with decreasing PUFA towards higher temperatures (Zeis et al., 2019). Masclaux et al. (2009) argued that unsaturation level of fatty acids increases towards lower temperature in membrane lipids, which, on the other hand, makes the animal more susceptible to heat-induced oxidative stress (Zeis et al., 2019). Here we show that *Daphnia* acclimated at 18°C for >1 year can successfully cope with a +6°C water temperature increase and counteract energy and PUFA limitation by *Planktothrix* if chytrids are also available in the diet. Our results suggest that trophic upgrading within the mycoloop is potentially a crucial and strong enough mechanism to support *Daphnia* fitness under a significant water temperature increase; e.g., during heatwaves. Extreme temperatures may amplify the deleterious effect of warming on the performance and fitness of ectothermic organisms (Kingslover et al., 2013), especially under reduced diet availability (Huey and Kingsolver, 2019). This certainly holds true for late summer phytoplankton in the temperate region, where large-sized hardly palatable phytoplankton may dominate in case of eutrophic conditions (Sommer et al., 1986; Ptacnik et al., 2008; O’Neil et al., 2012). Although our data suggest a positive diet quality effect of trophic upgrading by chytrids between a cyanobacterium and *Daphnia*, cyanobacteria lack growth-supporting sterols (Martin-Creuzburg et al., 2010) and thus trophic upgrading is limited to somatic effects of fatty acids for *Daphnia* and, importantly, subsequent consumers. Since we lack experimental evidence for the combined diet quality effect of sterols and PUFA-upgrading at the phytoplankton-zooplankton interface including chytrids, here we speculate that trophic upgrading by chytrids is a valid mechanism for supporting *Daphnia* performance under heat, irrespective of the type of algae (Rasconi et al., 2020). The elemental composition of phytoplankton can also constrain zooplankton fitness in particular under low phosphorus concentrations (Elser et al., 2001). *Daphnia* feeding on chytrids, however, remained unchanged in their C:P ratios, irrespective of water temperature, even though the chytrid zoospores contained significantly more phosphorus than *Planktothrix* and the chytrid-infected *Planktothrix* (Fig. S2). Rather, lipids and their FA, such as EPA and ARA, were the key molecules that could improve *Daphnia* fitness (Müller-Navarra et al., 2000; Brett et al. 2009). While Daphnia can adapt EPA to higher water temperature to sustain membrane fluidity if diet is neither limiting in quantity nor quality (Schlechtriem et al., 2006), they may function differently in case of highly limiting diet conditions. We argue that Daphnia preferentially bioconvert along with n-3 PUFA ALA to EPA to sustine first survival, then growth and reproduction, a process unaffected by a +6°C water temperature increase. While zooplankton fitness decreases in response to the dietary lack of ARA and EPA (Gulati and Demott, 1997), here we show that chytrids can support *Daphnia* fitness via sustained endogenous EPA and ARA conversion, even under heat. As global warming further intensifies deleterious algal blooms (Paerl and Huisman, 2008; Wagner and Adrian, 2009), the alternative energy and PUFA transfer via the mycoloop may become an increasingly crucial process to ensure pelagic food web functioning.

## Conclusions

Here we show that the presence of chytrids in the diet of the keystone herbivore, *Daphnia*, supports its fitness under increased water temperature, resulting in sustained survival, growth and successful reproduction. Chytrids support and improve PUFA accrual in *Daphnia* via increased dietary access to the n-3 PUFA SDA (preferential n-3 upgrading by the chytrids), which co-occurs with higher n-3 (ALA to EPA) than n-6 (LIN to ARA) PUFA upgrading by the *Daphnia* itself. We observed that the chytrid-synthesised SDA, which is retained selectively by *Daphnia*, remained quantitatively unchanged with heat. The heat significantly increased the efficiency of LIN to ARA bioconversion, with yet unknown benefits for the consumer. We conclude that the positive dietary effect of chytrids in terms of PUFA is a crucial mechanism at the phytoplankton-zooplankton interface, potentially further improving PUFA availability for higher trophic levels and so stable ecosystem functioning in aquatic ecosystem under global warming.

## Acknowledgement

We thank Samuel-Karl Kämmer and Katharina Winter, WasserCluster Lunz, for their help with preparations for the experiment and laboratory analyses. We thank Dr Thomas Rohrlack for providing the Planktothrix and chytrid isolates.

## Authors contributions

Conceptualisation: SR, MJK, RP. Developing methods: SR, AA, MJK, RP. Conducting the research: AA. Data analysis. AA. Data interpretation: AA, MP, MJK. Preparation figures & tables: AA. Writing: AA wrote the original draft and then all authors edited and commented on the drafts.

## Data availability statement

The data of this study are available from the corresponding author upon reasonable request.

## Supplementary materials

### Survival, growth and egg production rates

**S1.**
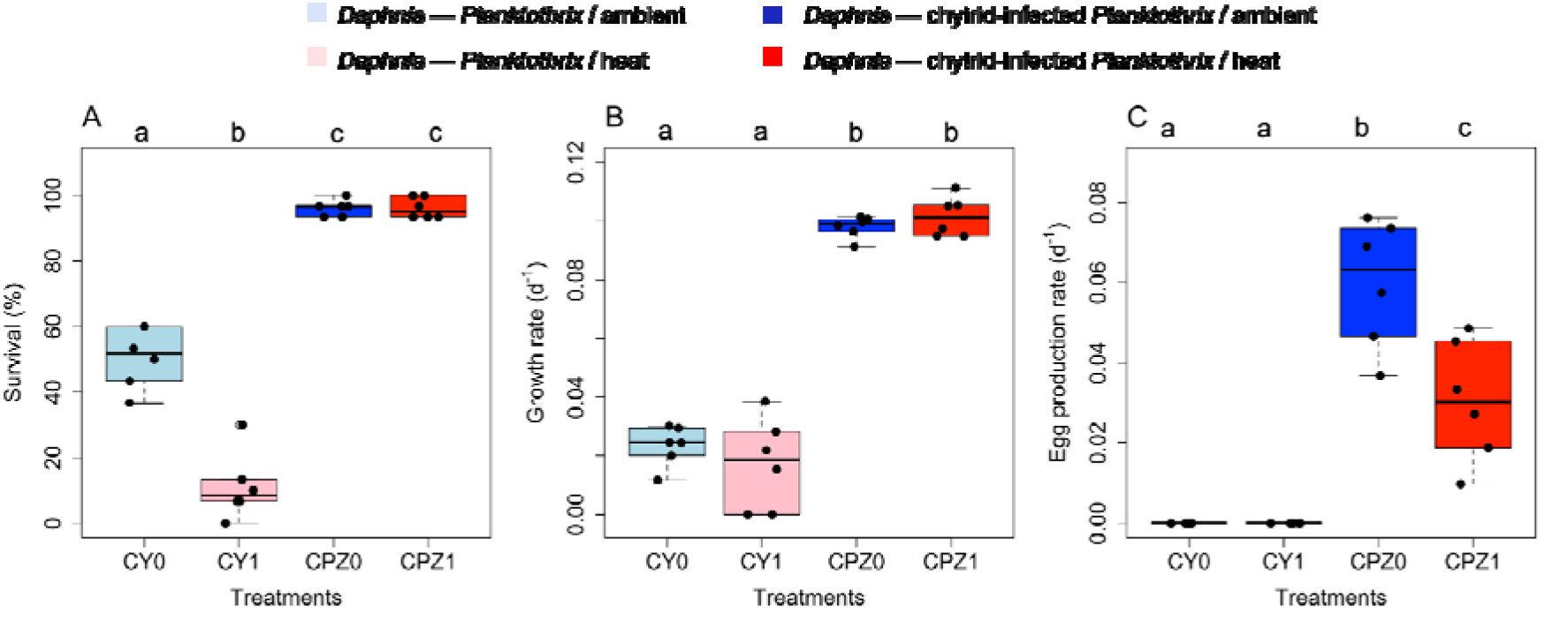
Boxplot of *Daphnia* **(A)** survival (% from 30 individuals per bottle) at day 11 (end of the heat treatment); **(B)** somatic growth rate; and **(C)** egg production rate among treatments. Letters above the boxplots denote the level of significance based on pairwise comparisons using Wilcoxon rank sum test (n=6 in all cases).

#### *Elemental composition of diets and* Daphnia

Carbon (C) and nitrogen (N) contents did not differ significantly among diets (Kruskal Wallis, df=2, p>0.05 in both cases). Phosphorus (P) differed significantly among diet sources (Kruskal-Wallis, chi-squared = 35.966, df = 2, p<0.001). It was significantly higher in chytrid zoospores compared with both *Planktothrix* and chytrid-infected *Planktothrix* (Pairwise Wilcoxon rank sum tests, p<0.001 in both cases). P in Planktothrix and chytrid-infected Planktothrix, however, did not differ significantly (Pairwise Wilcoxon rank sum test, p>0.05). The C:N ratio (Kruskal-Wallis, chi-squared = 22.92, df = 2, p<0.001) and the C:P ratio significantly differed among diet treatments (Kruskal-Wallis, chi-squared = 33.479, df = 2, p<0.001).

Chytrid zoospores were significantly lower in C:N (p_CY-Z_<0.001, p_CP-Z_<0.01) and C:P (p_CY- Z_<0.001, p_CP-Z_<0.001) compared with the other diet sources (S2A,B; Pairwise Wilcoxon rank sum test). Pairwise Wilcoxon rank sum test). The C:P ratio was also significantly lower in the chytrid zoospores compared with the cyanobacterium and the chytrid-infected cyanobacterium (p<0.001 in both cases; Supplement Fig S2B, Pairwise Wilcoxon rank sum test).

**S2.**
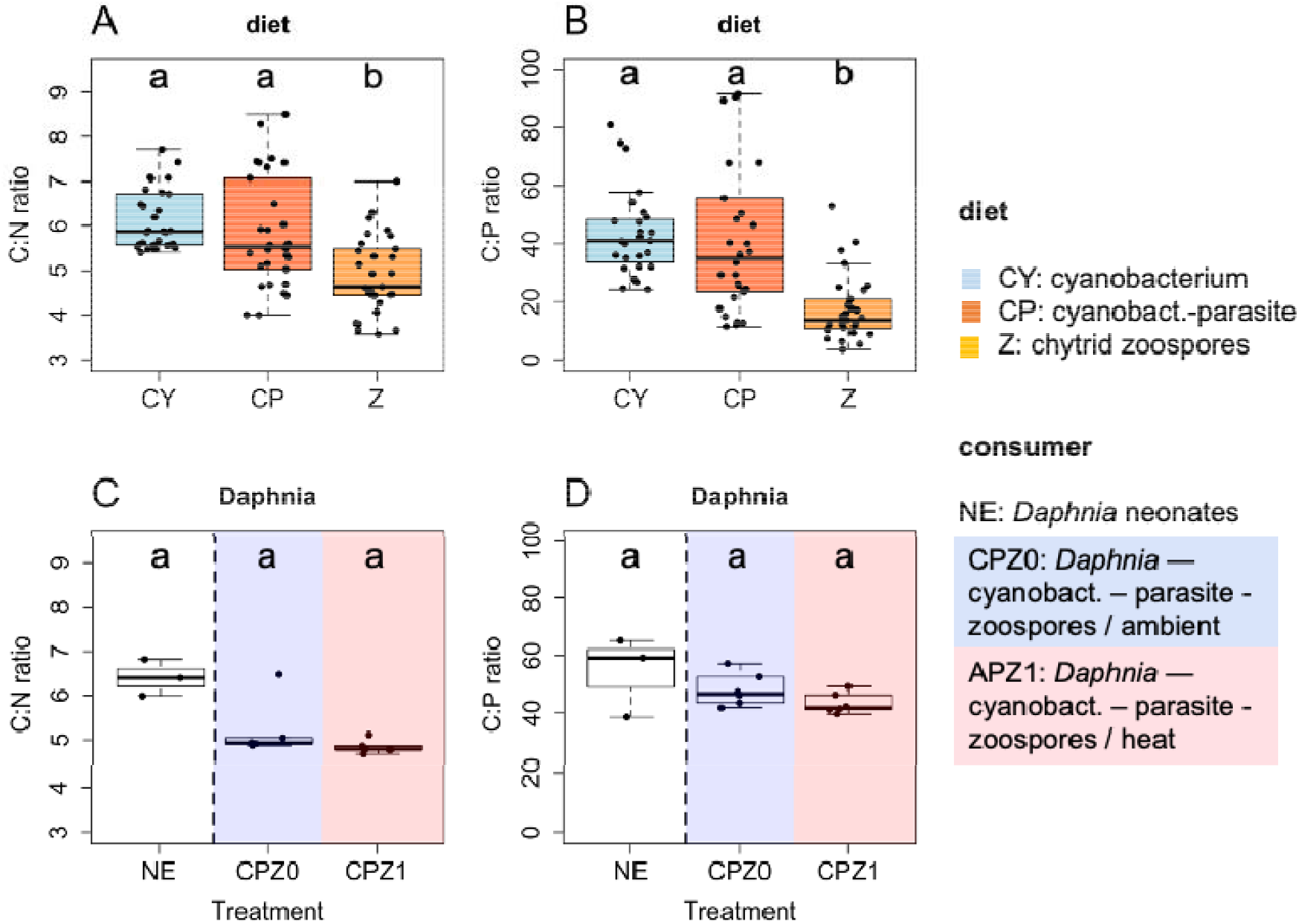
Boxplots of atomic **(A)** C:N and **(B)** C:P ratios in the diets(n=30 for each diet source); and boxplots of the atomic **(C)** C:N and **(D)** C:P ratios in *Daphnia* (n=3 for *Daphnia* neonates, and n=6 for *Daphnia* at the onset of the first successful reproduction, respectively). Letters above the boxplots denote the level of significance based on pairwise comparisons using Wilcoxon rank sum test.

The C and N content of *Daphnia* differed significantly among neonates and *Daphnia* feeding on chytrid-infected (Kruskal-Wallis rank sum test, df=2, p<0.01 in both cases). *Daphnia* feeding on the chytrid-infected diet had significantly higher C and N contents at the ambient compared with the heat treatment (Pairwise Wilcoxon rank sum test, p<0.01). The P content, however, did not differ significantly among neonates and *Daphnia* under different water temperature treatments (Pairwise Wilcoxon rank sum test, p>0.05).

The C:N ratio in *Daphnia* weakly but significantly differed among treatments (Kruskal-Wallis, chi-squared = 8.6, df = 2, p<0.05). However, in the pairwise comparisons, there was no significant differences among the *Daphnia* neonates and treatments (Pairwise comparisons using Wilcoxon rank sum test, p>0.05 in all cases). The C:P ratio in *Daphnia did not* differed significantly among treatments (Kruskal-Wallis, chi-squared = 2.4167, df = 2, p>0.05).

#### *PUFA profile of diets and* Daphnia

The key PUFA of diets were LIN (18:2n-6), ALA (18:3n-3) and SDA (18:4n-3; S3A). The *Planktothrix* and chytrid-infected *Planktothrix* did not differ in LIN and ALA (Pairwise comparisons using Wilcoxon rank sum test, p>0.05), but chytrid zoospores had significantly lower LIN and ALA compared with the cyanobacterium and its chytrid-infected cultures (Pairwise comparisons using Wilcoxon rank sum test, p<0.001 in both cases). The sole *Planktothrix* culture did not contain SDA, while the chytrid-infected *Planktothrix* and chytrid zoospores did contain SDA in similar amounts (Pairwise comparisons using Wilcoxon rank sum test, p_CP-Z_>0.05, p_CY-CP_<0.001, p_CY-Z_<0.001).

**S3.**
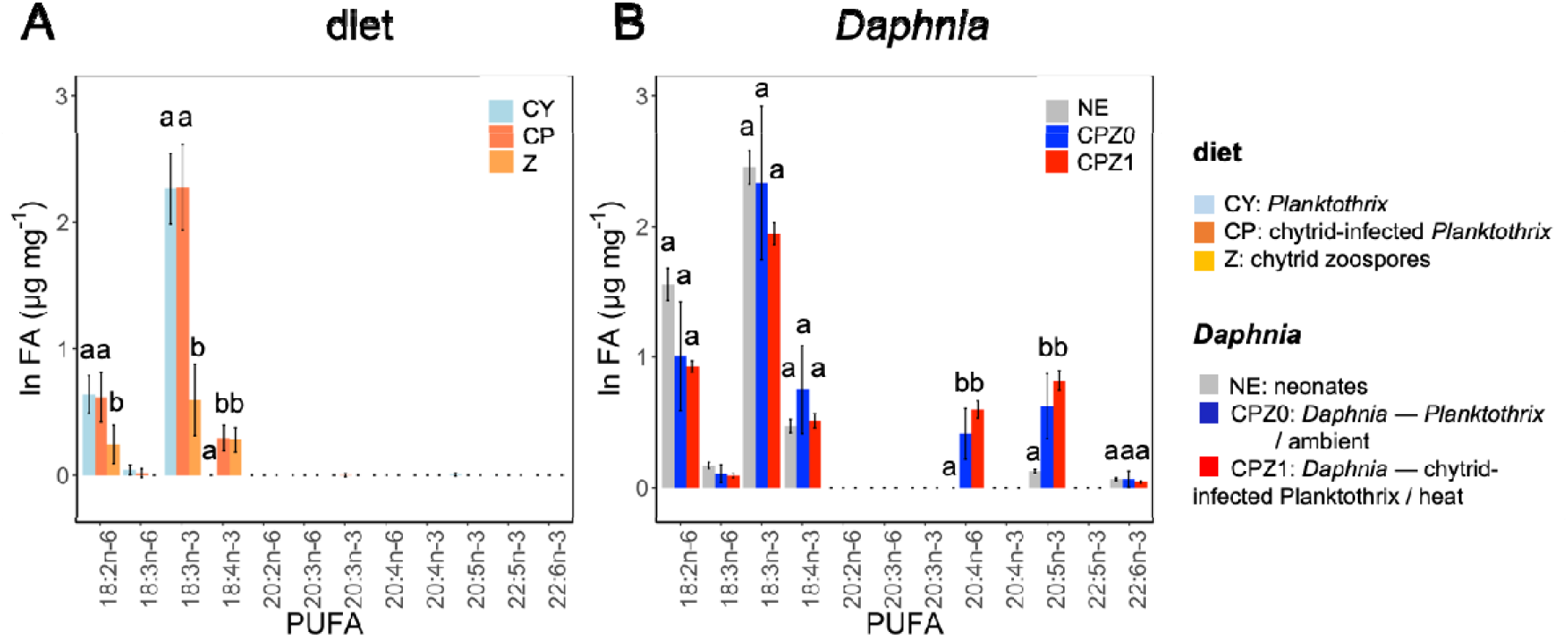
**(A)** Polyunsaturated fatty acid (PUFA) content of diets (n=15); and **(B)** PUFA content of *Daphnia* (n=3 for *Daphnia* neonates, and n=6 for *Daphnia* at the onset of the first successful reproduction, respectively). Letters denote the level of significance based on pairwise comparisons using Wilcoxon rank sum tests for each PUFA.

*Daphnia* neonates and *Daphnia* feeding on chytrid-infected diet did not differ significantly in LIN, ALA and SDA, irrespective of water temperature treatments (Pairwise comparisons using Wilcoxon rank sum test, p>0.05 in all pairwise cases). *Daphnia* feeding on chytrids, however, contained significantly higher amount of ARA (20:4n-6)and EPA (20:5n-3) compared with *Daphnia* neonates, irrespective of water temperature treatments (Pairwise comparisons using Wilcoxon rank sum test, p<0.05 in all pairwise cases).

